# Why are children so distractible? Development of attention and motor control from childhood to adulthood

**DOI:** 10.1101/747527

**Authors:** R. S. Hoyer, H. Elshafei, J. Hemmerlin, R. Bouet, A. Bidet-Caulet

## Abstract

Distractibility is the propensity to behaviorally react to irrelevant information. Though children are more distractible the younger they are, the precise contribution of attentional and motor components to distractibility and their developmental trajectories have not been characterized yet. We used a new behavioral paradigm to identify the developmental dynamics of components contributing to distractibility in a large cohort of French participants balanced, between age groups, in gender and socio-economic status (N=352; age: 6-25). Results reveal that each measure of these components, namely voluntary attention, distraction, impulsivity and motor control, present a distinct maturational timeline. In young children, increased distractibility is mostly the result of reduced sustained attention capacities and enhanced distraction, while in teenagers, it is the result of decreased motor control and increased impulsivity.

## Introduction

Remember a time when you were at school, listening to your teacher, and a driver honking their horn in the street or a classmate laughing might have caught your attention. These distractors interrupted your listening and note-taking. Healthy adults can easily focus on the task at hand again, unless the task-irrelevant distractor is significant or vitally important and requires changing behavior (e.g. a fire alarm). This capacity to be both task-efficient and aware of the surroundings without being constantly distracted requires a balance between voluntary and involuntary forms of attention. Voluntary attention enables performing an ongoing task efficiently over time by selecting relevant information and inhibiting irrelevant stimuli, while involuntary attention is captured by an unexpected salient stimulus, leading to distraction. Distractibility can then be considered as an attention state determining the propensity to have attention captured by irrelevant information and to behaviorally react to this information. At the brain level, voluntary and involuntary attention rely on partially overlapping dorsal and ventral brain networks, respectively (Corbetta & Shulman, 2002). In adults, the dorsal and ventral networks of attention have been found to be supported by activity in different frequency bands (alpha and gamma, respectively) and to interact within the prefrontal cortex (ElShafei, Bouet, et al., 2018; ElShafei et al., 2019). Increased distractibility with healthy aging has been associated with an imbalance between voluntary and involuntary attention spearheaded in the prefrontal cortex (Elshafei et al., 2020). Behaviorally, children are more distractible (Wetzel et al., 2015) compared to adults, though the interplay between voluntary and involuntary attention has been poorly documented in children. In ecological environments that are rich in distracting information, increased distractibility can be caused by (i) a reduced capacity to voluntarily pay attention to relevant events, (ii) an enhanced reaction to unexpected irrelevant distractors, or (iii) both. A better understanding of the causes of increased distractibility is crucial to improve rehabilitation or training programs to boost attention.

At the level of the brain, maturation progresses in a back-to-front direction from 6 to 18 years old (Toga et al., 2006). This cortical maturation appears to occur first in the primary sensory-motor cortex and progress into secondary, multimodal, and eventually supramodal cortices through childhood, with a protracted maturation of the prefrontal cortex (Gogtay et al., 2004; Sowell et al., 2004; Toga et al., 2006). Regarding attentional functioning, this postero-anterior maturation gradient is likely to support a shift in strategy from a posterior involuntary orienting network to engagement of an anterior voluntary attention network (Padilla et al., 2014; Smith et al., 2011), and a shift from a reactive to a proactive cognitive control (Andrews-Hanna et al., 2011; Chevalier et al., 2014; Lucenet & Blaye, 2014; Munakata et al., 2012).

Among the different facets of voluntary attention, two components related to distractibility have been behaviorally investigated: attentional orienting and sustained attention (Kanaka et al., 2008; Lin et al., 1999; Posner, 1980, 2012). The processing of relevant information can be proactively amplified by voluntary attentional processes allowing improved anticipation of upcoming events and sustained focus on ongoing tasks (Doebel et al., 2017; Gaspar & McDonald, 2014; Marini et al., 2016; Sawaki et al., 2012; Serences et al., 2004). Voluntary orienting of attention operates by proactively enhancing the processing of relevant information and inhibiting irrelevant events (van Zomeren & Brouwer, 1994; Posner, 1980, 2012). Posner paradigms with endogenous informative or uninformative cues have been used to measure the voluntary orienting of attention in anticipation of a target in children (Mezzacappa, 2004; Reis Lellis et al., 2013; Ross-Sheehy et al., 2015; Rueda et al., 2004). Results are conflicting: some show that the capacity to voluntarily orient attention is mature in children (Rueda et al., 2004) and even before the age of six (Ross-Sheehy et al., 2015), while others show that the benefit in reaction times (RT) to targets following informative cues increases from the age of 6 to adulthood (Mezzacappa, 2004; Reis Lellis et al., 2013).

These findings suggest that the voluntary orienting of attention may improve during childhood, but its precise developmental trajectory remains unclear. Sustained attention is the ability to maintain the attentional focus over time on a given task (Betts et al., 2006; Oken et al., 2006; Parasuraman et al., 1989). It relies on tonic arousal, also called vigilance (Levy, 1980). In children, sustained attention was mostly measured using detection tasks of targets among non-target stimuli presented at a fast rate (e.g. Continuous Performance Test; Conners & Sitarenios, 2011). A reduction in RT variability, as well as in the number of false alarms and missed responses, has been observed from the age of 5 until early adulthood (Betts et al., 2006; Kanaka et al., 2008; Lin et al., 1999; Thillay et al., 2015). These findings suggest a continuous maturation of sustained attention throughout childhood and adolescence with critical maturation steps at 6 and 13 years old. To our knowledge, no study has investigated the developmental trajectory of sustained attention in a more ecological context including distracting events.

Only a few studies have attempted to characterize the impact of distracting events on ongoing performance in children. Unexpected irrelevant salient sounds induce distraction (i.e. the reactive allocation of attention and resources to this salient event followed, if possible, by a reallocation of attention and resources towards the task), resulting in a behavioral cost in the task at hand (Bidet-Caulet et al., 2015; Escera et al., 2000; Näätänen, 1992). But they also trigger a burst of phasic arousal according to the sound properties, resulting in behavioral benefits (Bidet-Caulet et al., 2015; Masson & Bidet-Caulet, 2019; Max et al., 2015a; Näätänen, 1992; Wetzel et al., 2012). Distraction was mostly investigated using audio-visual oddball paradigms, involving the discrimination of targets preceded by task-irrelevant standard or novel sounds (Escera et al., 2000; Wetzel et al., 2012; Wetzel & Schröger, 2007, 2014). Lower hit rates and longer reaction times to targets preceded by novel sounds are considered a measure of deviance distraction. These measures were found to improve from childhood to adulthood (Olesen et al., 2007; Wetzel et al., 2006; Wetzel & Schröger, 2007), suggesting a reduction in distraction with age. The burst of arousal triggered by salient sounds would be mediated by the norepinephrine system, and it results in a transient and non-specific state of readiness to respond to any upcoming stimulus (Aston-Jones & Cohen, 2005; Corbetta et al., 2008). Phasic arousal to sounds can be indirectly measured through pupil dilation (Widmann et al., 2018) which was found contingent to brain maturation processes (Wetzel, Schröger, et al., 2016). Previous works have shown that an increase in phasic arousal can also lead to an increased false alarm rate (Aston-Jones & Cohen, 2005; Duncan et al., 2016). Impulsivity is the tendency to act without forethought and to fail to appreciate circumstances related to the present situation (Barratt & Patton, 1983; Stanford et al., 2009). An increased false alarm rate is typically observed in impulsive persons and could result from an enhanced phasic arousal (Eysenck & Eysenck, 1985; Houston & Stanford, 2001; Zhang et al., 2015) coupled – or not – with a lack in motor control (Booth et al., 2003; Kanaka et al., 2008; van den Wildenberg & Crone, 2005; Wright et al., 2003). False alarms in the absence of a distractor might reflect difficulty in maintaining an efficient proactive control. By contrast, a false alarm following the occurrence of a distractor, or a response to the distractor itself, might indicate a difficulty in reactive control. Distracters can indeed be suppressed reactively, when a salient but irrelevant stimulus captures attention and its behavioral consequence needs to be wiped out (Gaspar & McDonald, 2014; Geng & DiQuattro, 2010). To that extent, a better understanding of the developmental trajectories of distraction, phasic arousal and impulsivity triggered by an unexpected salient event would help disentangle the maturation of proactive and reactive attentional control.

In sum, previous behavioral studies have shown that voluntary orienting of attention, sustained attention, distraction, phasic arousal, impulsivity and motor control follow different developmental trajectories, which remain to be specified. Despite the importance of distractibility, its developmental trajectory is currently unknown. The aim of the present study is to specify the maturational timeline of the different components of distractibility in people between 6 and 25 years old. We used an adaptation of a recently developed paradigm, the Competitive Attention Task (CAT; Bidet-Caulet et al., 2015). This paradigm combines the Posner task and the oddball paradigm to provide simultaneous and dissociated measures of voluntary attention, distraction, phasic arousal, impulsivity and motor control. To assess voluntary attention orienting, the CAT includes informative and uninformative visual cues respectively indicating – or not – the spatial location of a forthcoming auditory target which is to be detected. To measure distraction, the CAT comprises trials with a task-irrelevant distracting sound preceding the target according to several delays. This change in distractor timing onset allows the dissociation of the effects of distraction and phasic arousal. Moreover, similarly to other detection tasks, the rates of different types of false alarms, late and missed responses provide measures of sustained attention, impulsivity and motor control.

The nature of the present study is rather exploratory; we nevertheless had several assumptions detailed in the followings and we used quite large population samples to test these hypotheses. Based on the literature on the development of both the brain and attention capacities, we hypothesized a move from a reactive to a proactive strategy for attentional control from early childhood to adulthood (Blackwell & Munakata, 2014; Chevalier et al., 2014; Munakata et al., 2012). More precisely, young children are expected to present stronger behavioral manifestations in reaction to distracting sounds such as increased rates of responses to distractors, but also of late and missed responses to targets following a distractor. Proactive attentional control is likely to mature later. Better voluntary attention performances (e.g. reduced RT variability, reduced late response and false alarm rates and larger cue effect on RTs in trials with no distractor) are expected during adolescence. In summary, distraction would first decrease during early childhood, followed by an improvement in motor control, and a protracted maturation of voluntary attention. As previous studies show an increased motor speed during spatial attention tasks in men, gender was taken into account in the present study (Gur & Gur, 2017; Jain et al., 2015; Merritt et al., 2005).

## Materials and Methods

### Participants

415 subjects of French nationality, from diverse socioeconomic statuses, participated in the study which took place from September 2016 to January 2018, in France, in large cities (Lyon and Grenoble) and in rural areas (Ardèche). Typically developing children from the 1st to 12th grade were recruited in public and private schools. Adults were recruited through flyers and email lists. Data from 63 participants were excluded from the analysis, either because of neurological disorders or substance use (N=9), auditory problems (N=2), non-compliance with the instructions (N=17), correct trial percentage < 50% in NoDis condition (N=11) or technical issues (N=24). A total of 352 subjects (88% right-handed, 51% female, 6 to 25 years old) were included in the analysis. All subjects had normal hearing and normal or corrected-to-normal vision. Participants were divided into 14 age groups (Table 1). For participants under the age of 18, signed informed consent was obtained from both parents, and assent was given by the children. All adult participants (18-25 years old) gave written informed consent. This study was approved by participating schools and was conducted according to the Helsinki Declaration, Convention of the Council of Europe on Human Rights and Biomedicine, and the experimental paradigm was approved by the French ethics committee Comité de Protection de Personnes for testing in adults and children.

**Table 1.**
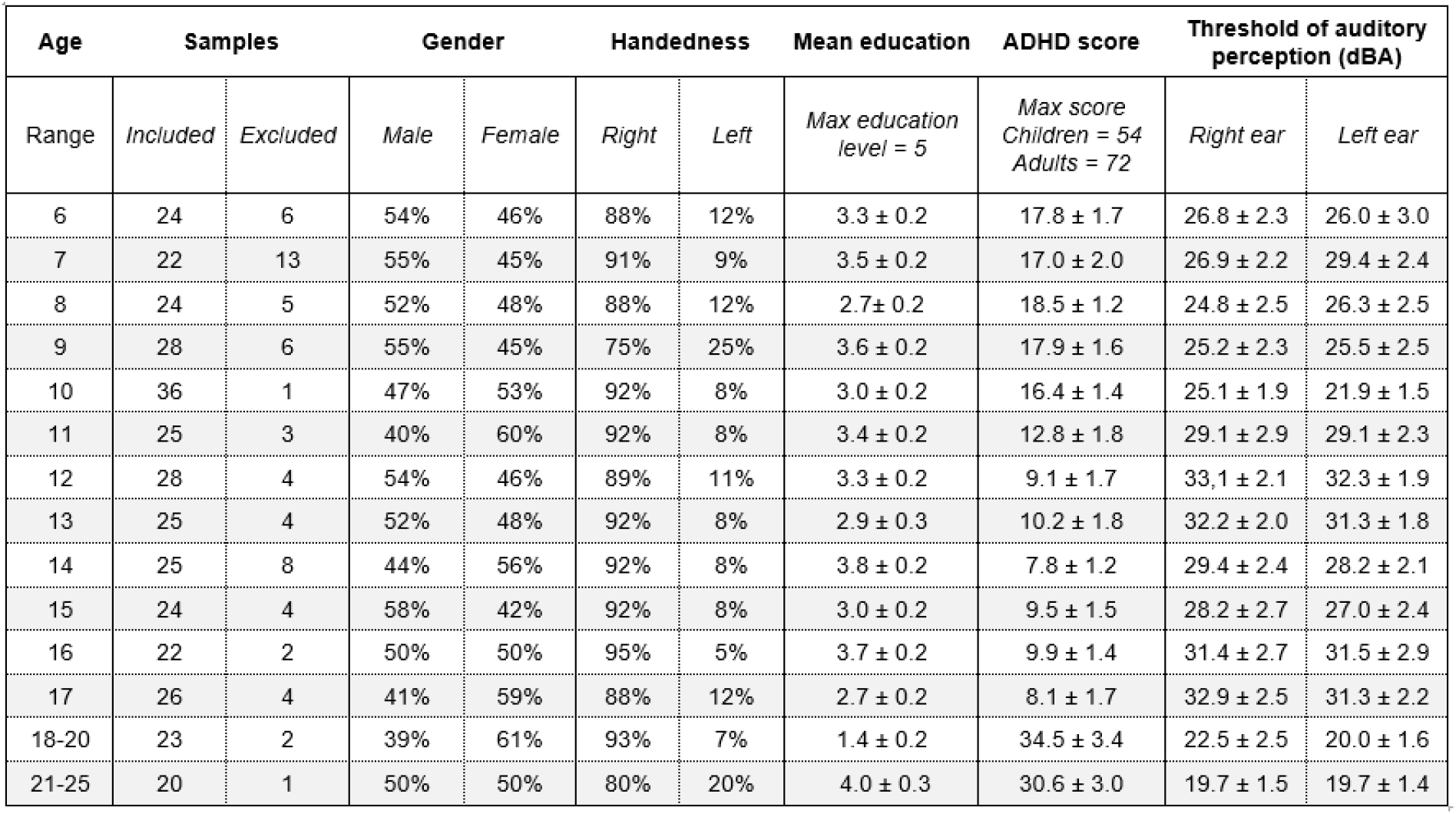
Characteristics of the population sample. *Note*. Detailed samples by age for gender, handedness, mean parent education level for children and mean education level for adults, total ADHD scale scores and thresholds of auditory perception (± standard error of the mean, SEM). Please note that the mean age in years and months in each group cannot be provided since only participants’ age in years was collected.

Groups were matched for gender (male or female) and handedness. Age groups from 6 to 17 years of age were matched for economic status (SES, see Fig. S1) and educational level of their parents (0 = no diploma, 1 = vocational certificate obtained after the 9^th^ grade, 2 = high school diploma; 3 = 12^th^ grade / associate’s degree; 4 = bachelor degree; 5 = master degree and further). The 18 to 25-year-old participants reported their own SES and education level: around 80% were students at the university and 20% were employees.

### Stimuli and Task

50% of the trials (Fig. 1a) consisted of a visual cue (200-ms duration) followed, after a 950-ms delay, by a 200-ms target sound. The cue was presented centrally on a screen with a gray background and could either be a dog facing left or right, or a dog facing forward. The target sound was the sound of a dog barking monaurally presented at 15 dB SL (around 43 dBA) in headphones.

**Figure 1.**
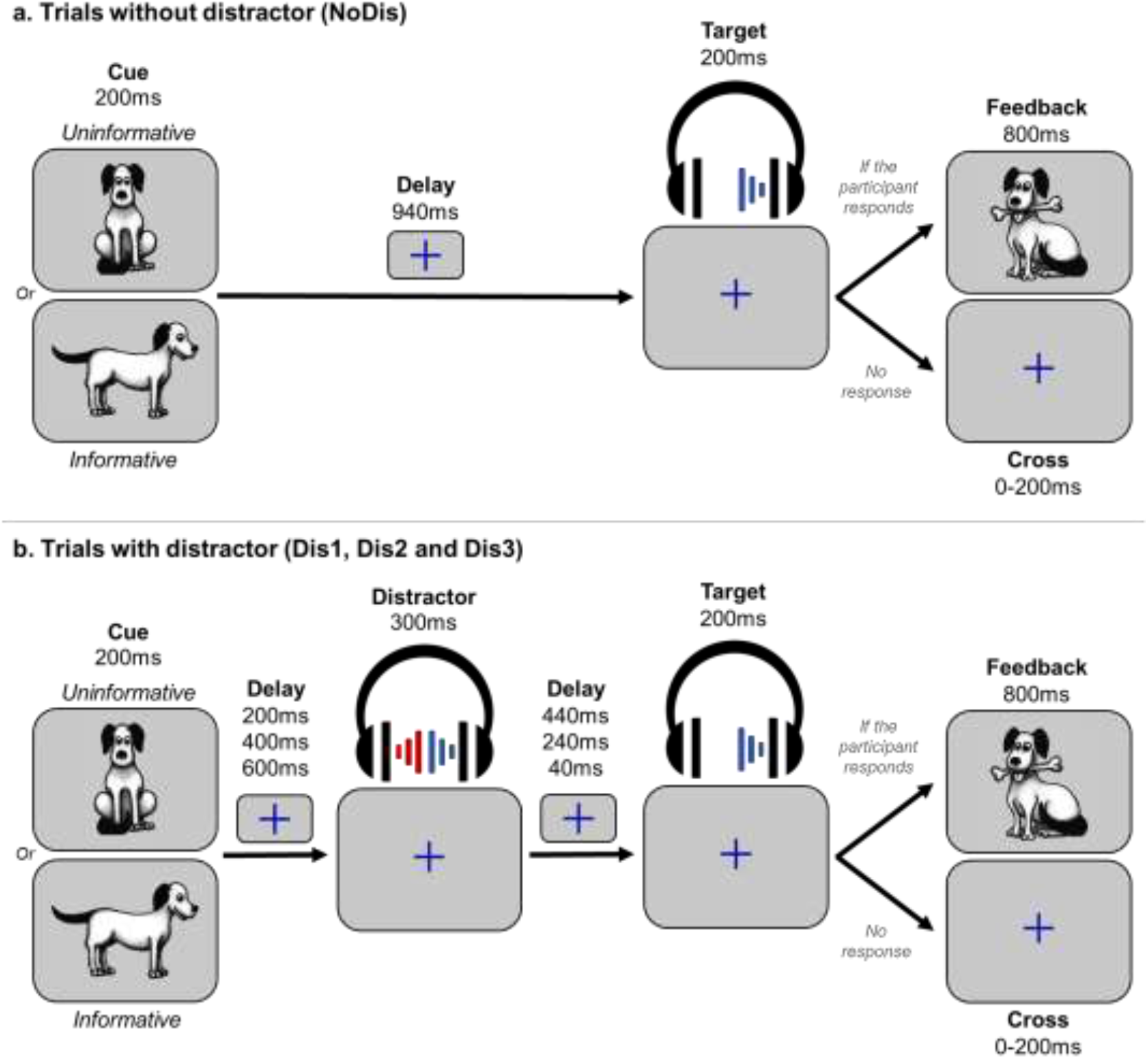
Protocol. *Note*. a) In uninformative trials, a facing-front dog was used as visual cue (200 ms duration), indicating that the target sound will be played in either the left or right ear. In informative trials, a facing left or right dog visual cue (200 ms duration) indicated in which ear (left or right, respectively) the target sound will be played (200 ms duration) after a delay (940 ms). If the participant gave a correct answer, a feedback (800ms duration) was displayed. b) In trials with distractor the task was similar, but a binaural distracting sound (300 ms duration) - such as a phone ring - was played during the delay between cue and target. The distracting sound could equiprobably onset at three different times: 200 ms, 400 ms, or 600 ms after the cue offset.

In the other 50% of the trials, the trial structure was identical, but a binaural distracting sound (300-ms duration) was played during the delay (Fig. 1b) at 35 dB SL (around 61 dBA). A total of 18 different distracting sounds were used (phone ringtone, clock-alarm, etc.) for each participant. The distracting sound could be equiprobably played at three different times during the delay: 200 ms (Dis1), 400 ms (Dis2) and 600 ms (Dis3) after cue offset. One block of the task contained 24 NoDis, 8 Dis1, 8 Dis2 and 8 Dis3 trials. Therefore, during the whole experiment (3 blocks), 72 NoDis, 24 Dis1, 24 Dis2 and 24 Dis3 trials were presented.

The proportion of cue and target categories were distributed equiprobably between trials with and without distracting sound. The informative condition represented 75% of the trials: in that case the dog was facing left or right, indicating the ear of the target sound presentation (37.5% left and 37.5% right). The uninformative condition represented 25% of the trials: the facing-front dog was followed by the target sound in the left (12.5%) or right (12.5%) ear.

To compare behavioral responses to acoustically matched sounds, the same distracting sounds were played for each distractor condition (Dis1, Dis2 or Dis3) in the informative condition. Each distracting sound was played 4 times during the whole experiment, but no more than twice during each single block to limit habituation.

Subjects were instructed to perform a detection task by pressing a key as fast as possible when they heard the target sound. They were asked to focus their attention to the cued side in the case of informative cue. Participants were informed that informative cues were 100% predictive and that a distracting sound could be sometimes played. In the absence of the visual cue, a blue fixation cross was presented at the center of the screen. Subjects were instructed to keep their eyes fixated on the cross (see Appendix S1 for full instructions). The training block was composed of 10 trials and was built following the same time course than the task. Two trials contained distractors which belonged to the pool of 18 distracting sounds used in the experiment.

When participants answered within 3300 ms after the target onset, a dog holding a bone (800-ms duration) was presented 500 ms after the response followed by the fixation cross for a randomized period of 1700 ms to 1900 ms. If the participant did not respond during the 3300 ms post target onset, the fixation cross was displayed on the screen for an additional randomized delay of 100 ms to 300 ms. At the end of each block, the average reaction time was indicated to the participant to motivate him/her to respond as fast as possible.

### Procedure

Participants were tested in small groups of 2 to 4. Adults were tested in the lab or at the university, and children were tested at school, all in a quiet room. Participants were seated in front of a laptop (approximately 50 cm from the screen) delivering pictures and sounds, as well as recording behavioral responses using Presentation software (Neurobehavioral Systems, Albany, CA, USA). Auditory stimuli were played through headphones.

First, the auditory threshold was determined for the target sound, in each ear, for each participant using the Bekesy tracking method. This resulted in an average target threshold across subjects of 28 ± 0.6 dBA (see Table 1 for details by age range). Then, participants performed a short training of the task followed by three 4-min blocks of 48 pseudo-randomize trials each. The order of the 3 blocks was randomized through participants using a Latin square. The experimenter gave verbal instructions to the children before the test. An experimental session lasted around 30 minutes. At the end of every experimental session, the experimenter explained the aim of the study to participants and took time to answer questions.

Adults and parents of children enrolled in the study filled out a short questionnaire about their SES characteristics and respectively completed the Adult Self-Report Scale (ASRS; Kessler et al., 2005) and the Attention-Deficit Hyperactivity Disorder Rating Scale IV (ADHD RS; DuPaul et al., 1998) questionnaires, both assessing symptoms of ADHD in adults and children according to the diagnostic criteria of the Diagnostic and Statistical Manual of Mental Disorders (Diagnostic and Statistical Manual of Mental Disorders, 2013). Adults also filled out the State-Trait Anxiety Inventory (STAI) Y-A and B (Spielberger et al., 1970) to evaluate anxiety as a state and trait. At the end, every minor participant answered a short post-experiment questionnaire about their motivation level, their focus state and stress level during the CAT. The aforementioned questionnaires were administered in order to collect normative data for other research projects and the analysis of the collected responses is beyond the scope of the present study.

### Measurement parameters

We used a custom MATLAB program to extract and preprocess behavioral data. First, we visually inspected the reaction time distribution relative to target onset for each age group (see Fig. S2). RT limits for correct responses were computed to consider the overall brain maturation effect on speed processing. For each participant, the longest reaction time for a correct response (RT upper limit) was calculated from all RT > 0ms using the John Tukey’s method of leveraging the Interquartile Range. The shortest reaction time for a correct response (RT lower limit) was calculated for each age range (see Fig. S2 and Appendix S2). Thus, (i) responses before the RT lower limit were considered as a part of the false alarm response type; (ii) responses between the lower and the upper RT limits were considered as correct responses; (iii) responses after the RT upper limits were considered as late responses.

To investigate the diversity of behaviors observed during the task, the following 8 behavioral measures were analyzed further (see Fig. S3):

- Positive reaction times (RT+, ms).
- RT standard deviation (RT SD, ms): mean standard deviation of RT+ in the NoDis condition for each block separately.
- Late response (LateRep, %): the percentage of responses performed in the NoDis condition during the period starting from the RT upper-limit to 3300 ms.
- Missed response (MissRep, %): the percentage of trials without any response made during the entire trial duration up to 3300 ms post-target.
- Cue response (CueRep, %): the percentage of responses performed during the 150-450ms period post-cue onset.
- Distractor response (DisRep, %): the percentage of responses performed during the 150-450 ms period post-distractor onset.
- Anticipated response (AntRep, %): the percentage of responses performed:

○ in NoDis and Dis1: from 300 ms pre-target to the RT lower limit post-target;
○ in Dis2: from 150 ms pre-target to the RT lower limit post-target;
○ in Dis3: from 100 ms post-target to the RT lower limit post-target.
- Random responses (RandRep, %): the percentage of responses performed in the remaining periods of the trials, i.e., within the 150 ms post-cue onset and:

○ in NoDis: during the 450 to 850 ms period post-cue onset;
○ in Dis1: during the 450 to 550 ms period post-cue onset;
○ in Dis2: during the 450 to 750 ms period post-cue onset;
○ in Dis3: during the 450 to 950 ms period post-cue onset.

### Statistical analyses

In order to test the similarity between measures or to estimate a degree of logical support or belief regarding specific results, Bayesian statistics were used. To estimate physical tendencies linked to the behavioral measurements, we used generalized linear mixed models (GLMM) in a frequentist approach.

### Socio-economic data analysis

To confirm that our sample population was similarly distributed across age ranges in block order, handedness, and gender, we performed Bayesian contingency table tests. For children from 6 to 17-years-old only, similar analysis was performed on SES and education level of the parents. We reported Bayes Factor (BF_10_) as a relative measure of evidence. To interpret the strength of evidence in favor of the null model (uniform distribution), we considered a BF between 0.33 and 1 as weak evidence, a BF between 0.1 and 0.33 as positive evidence, a BF between 0.01 and 0.1 as strong evidence and a BF lower than 0.01 as a decisive evidence. Similarly, to interpret the strength of evidence against the null model, we considered a BF between 1 and 3 as weak evidence, a BF between 3 and 10 as positive evidence, a BF between 10 and 100 as strong evidence and a BF higher than 100 as a decisive evidence (Lee & Wagenmakers, 2013).

Statistical analyses of hearing threshold are presented in Table S1 and Appendix S3. Statistical analyses of attention scores are presented in Appendix S4.

Bayesian statistics were performed using JASP^®^ software (JASP Team (2018), JASP (Version 0.9) [Computer software]).

### Behavioral data analysis

#### Frequentist statistical approach

We expected a large between-individual variability in RT+ and response type rates according to conditions. This behavioral response variability limits the comparison of data between the conditions and means that data cannot simply be pooled for analysis. GLMM are the best way to deal with such datasets, as they allow for correction of systematic variability (Bates et al., 2015). We accounted for the heterogeneity of performances between-subjects and experimental conditions by defining them as effects with a random intercept and slopes, thus instructing the model to correct for any systematic differences in variability between the subjects (between-individual variability) and condition (between-condition variability). To confirm the need for mixed nested models, we used a likelihood ratio analysis to test the model fit before and after sequential addition of random effects. We used the Akaike Information Criterion and the Bayesian Information Criterion as estimators of the quality of the statistical models generated (Matuschek et al., 2017). To optimize our model, we checked the normality of the model residual.

To assess the impact of the manipulated task parameters (cue information and distractor type) and of participant demographics (age and gender), on each type of behavioral measure (RT, RT SD, LateRep, MissRep, CueRep, DisRep, AntRep, RandRep), we analyzed the influence of four possible fixed effects (unless specified otherwise in the next section):

1. between-subject factor AGE: 14 levels (see Table 1);
2. between-subject factor GENDER: 2 levels (male and female);
3. within-subject factor CUE and CUELRN: 2 levels (CUE: informative vs. uninformative) for measures recorded after the target onset (Hit, LateRep and MissRep) and 3 levels (CUELRN: left, right and neutral) for the measures recorded before the target onset (CueRep, RandRep, DisRep and AntRep);
4. within-subject factor DISTRACTOR: 4 levels (NoDis, Dis1, Dis2 and Dis3), except for DisRep: 3 levels (Dis1, Dis2 and Dis3).

A summary of the data and factors used in statistical modeling can be found in Table 2. Please note that for response types cumulating less than 1 % of response proportion across the total sample (cue and random responses), only 2-way interactions were considered.

**Table 2.** Main statistical analyses according to behavioral response types. *Note*. Experimental conditions, factors and models used as a function of the behavioral measurement. *Response type cumulating less than 1% of response proportion across the total sample (only 2-way interactions were considered). Detailed factor levels: CUE = informative vs. uninformative; CUELRN = left, right and neutral; Block = first, second and third. Models: GLMM = Generalized Linear Mixed Model.

| Response types | Condition(s) used for response types calculation | Fixed factor(s) |  | Random factor(s) | Distribution fitting | Missing data |
| --- | --- | --- | --- | --- | --- | --- |
|  |  | Between subjects | Within subjects |  |  |  |
| RT (log) | NoDis vs. Dis1 vs. Dis2 vs. Dis3 | Age, Gender | Cue, Distractor | Distractor + Subject | Gaussian | 2.4 % |
| SD RT (log) | NoDis | Age, Gender | Block | Subject | Gaussian | 2.4 % |
| Late responses | NoDis | Age, Gender | Cue | Subject | Binomial | 0.0 % |
| Missed responses | NoDis vs. Dis1 vs. Dis2 vs. Dis3 | Age, Gender | Cue, Distractor | Subject | Binomial | 0.0 % |
| Cue responses * | NoDis & Dis1 & Dis2 & Dis3 | Age, Gender | CueLRN | Subject | Binomial | 0.0 % |
| Distractor responses | Dis1 & Dis2 & Dis3 | Age, Gender | CueLRN | Subject | Binomial | 0.0 % |
| Anticipated responses | NoDis vs. Dis1 | Age, Gender | CueLRN, Distractor | Distractor + Subject | Binomial | 0.0 % |
| Random responses * | NoDis vs. Dis1 vs. Dis2 vs. Dis3 | Age, Gender | CueLRN | Subject | Binomial | 0.0 % |

We ran a type II analysis of variance. Wald chi-square tests were used for fixed effects in linear mixed-effects models (Fox & Weisberg, 2018). We only considered the explanatory variables. The fixed effect represents the mean effect across all subjects after correction of variabilities. Frequentist models and statistics were performed in R^®^ 3.4.1 using the lme4 (Bates et al., 2015) and car (Fox & Weisberg, 2018) packages. We only considered results of main analyses significant at p < .01.

When we found a significant main effect or interaction - and did not plan ahead for specific post-hoc analysis - Post-hoc Honest Significant Difference (HSD) post-hoc tests were systematically performed using the emmeans package (emmeans version 1.3.2). P-values were considered as significant at p < .05 and were adjusted for the number of performed comparisons.

In the Results section, we reported the SEM as the estimator of the distribution dispersion of the measures of interest, when not specified.

##### Reaction Times

To be sure to perform analyses on a sufficient number of trials per condition, participants with less than 45 NoDis trials or than 6 trials in each of the distractor conditions with positive RT were excluded from RT+ analysis (leading to a total average of trials with positive RT of 68 ± 0.2, 19 ± 0.2, 20 ± 0.2 and 20 ± 0.2, in the NoDis, Dis1, Dis2, and Dis3 conditions, respectively). Based on visual inspection of median RT distribution in distractor conditions, one outlier participant was also identified and removed from this analysis. Revised samples for RT+ analysis are: 6-year-olds: n = 17; 7-year-olds: n = 20). The percentage of missing data over the total sample of included subjects in analyses is shown in Table 2.

Raw RT were log-transformed at the single trial scale to be better fitted to a linear model with Gaussian family distribution.

For post-hoc analysis of the DISTRACTOR by AGE interaction on RT+, we planned to analyze two specific measures of the distractor effect: the distractor occurrence (median RT+ in NoDis minus median RT+ in Dis1) and the distractor position (median RT+ in Dis3 minus median RT+ in Dis1). Based on previous results (Bidet-Caulet et al., 2015; Masson & Bidet-Caulet, 2019), these differences can be respectively considered as good approximations of the facilitation and detrimental distraction effects triggered by distracting sounds (see Fig. S4). We first performed Shapiro tests to estimate the normality of the arousal and distraction effects.

Planned non-parametric Kruskal-Wallis tests with the AGE as between-subject factor were applied to these measures when the data were not normally distributed. When the Kruskal-Wallis test revealed a significant effect of the AGE, we performed non-parametric paired Nemenyi post-hoc tests to identify developmental stages across age ranges.

Normality check and non-parametric statistics were performed in R^®^ 3.4.1 using the stats (R Core Team, 2017) and DescTools (Andri Signorell et mult. al., 2019) packages.

##### Other measures

RT SD were log-transformed at the single trial scale to be better fitted to a linear model with Gaussian family distribution, with the fixed factors AGE and GENDER as between-subject factor and BLOCK (3 levels) as within subject factor.

Response types were re fitted to a linear model with binomial distribution without transformation. GLMMs extend the linear mixed model to deal with non-normally distributed data. In particular, GLMMs can handle binary data. In that respect, GLMMs analysis consider a number of data points equal to the total number of trials presented during the test, in each subject.

LateRep were fitted to a linear model with binomial distribution without transformation, with fixed factors AGE and GENDER as between-subject factor and CUE (2 levels) as within subject factor. Only, the NoDis condition was analyzed since few participants committed LateRep in distractor conditions (total average: 3.5 ± 0.1%).

MissRep were fitted to a linear model with binomial distribution without transformation, with fixed factors AGE and GENDER as between-subject factor and CUE (2 levels) and DISTRACTOR (4 levels) as within subject factor.

CueRep and Randrep were fitted to a linear model with binomial distribution without transformation, with fixed factors AGE and GENDER as between-subject factor and CUELRN (3 levels) as within subject factor. As all participants made in average less than 1% of CueRep and RandRep, responses were considered across all DISTRACTOR levels (NoDis, Dis1, Dis2, Dis3) and their modelization was limited to two-way interactions.

DisRep were fitted to a linear model with binomial distribution without transformation, with fixed factors AGE and GENDER as between-subject factor and CUELRN (3 levels) as within subject factor. Responses were considered across DISTRACTOR levels (Dis1, Dis2, Dis3).

AntRep were fitted to a linear model with binomial distribution without transformation, with fixed factors AGE and GENDER as between-subject factor and CUELRN (3 levels) as within subject factor. Because of the important differences in the duration of the AntRep windows between distractor conditions (see Fig. S3), the GLMM was performed on the NoDis and Dis1 conditions, only (same time-frame for AntRep in these two conditions).

#### Planned Bayesian regressions

To confirm a potential age effect on specific measures, Planned Bayesian Kendall regressions with age were performed on specific RT+ normalized measures of attention:

- Voluntary attention orienting: (medianRT_+NoDisUninf_ – medianRT_+NoDisInf_) / medianRT_+AllConditions_;
- Arousal effect: (medianRT_+NoDis_ – medianRT_+Dis1_) / medianRT_+AllConditions_;
- Distraction effect: Voluntary attention orienting: (medianRT_+Dis3_ – medianRT_+Dis1_) / medianRT_+AllConditions_.

## Results

### Population characteristics

Using Bayesian contingency table tests, we found decisive evidence for a similar distribution in block order (BF_10_ = 2.1·10^-5^), gender (BF_10_ = 5.2·10^-7^) and handedness (BF_10_ = 8.1·10^-21^) across age ranges. We observed – in the 6 to 17-year-olds – a decisive evidence for a similar distribution in SES (BF_10_ = 8.9·10^-20^) and education level of the parents (BF_10_ = 1.5·10^-19^) across age ranges.

### Behavioral Data

We extracted 8 behavioral measures from participants’ responses: positive reaction times (RT+), RT+ standard deviation (RT SD) as a measure of sustained attention, late response % (LateRep) as a measure of attentional lapses, missed response % (MissRep) and distrastor response % (DisRep) as measures of distraction, cue response % (CueRep) and anticipated response % (AntRep) as measures of impulsivity, and random response % (RandRep) as a measure of motor control.

For each type of behavioral measurement, we analyzed the influence of AGE, GENDER, CUE and DISTRACTOR factors (unless specified otherwise in the Table 2). In the following, when a factor was involved in a main effect and a higher order interaction, we only specified the post-hoc analysis related to the interaction.

### RT+

RT+ were modulated by GENDER (χ^2^ (1) = 18.1; p < .001): male (325.6 ± 1.6 ms) were faster than female (350.8 ± 1.7 ms) participants.

We observed main effects of AGE (χ^2^ (13) = 460.0; p < .001, Fig. S5), CUE (χ^2^ (1) = 56.1; p < .001) and DISTRACTOR (χ^2^ (3) = 1326.5; p < .001). We did not observe a CUE by AGE interaction (Fig. 3a). This was confirmed by positive evidence against a correlation of the voluntary attention orienting normalized measure with age (Kendall’s Tau = 0.041, BF_10_ = 0.1).

A DISTRACTOR by CUE interaction was significant (χ^2^ (3) = 26.6; p < .001; Fig. 2). Post-hoc HSD tests showed that participants were faster to detect targets preceded by an informative cue in the NoDis, Dis2 and Dis3 (p <.001) conditions, while no cue effect was found in the Dis1 condition (p = .694).

**Figure 2.**
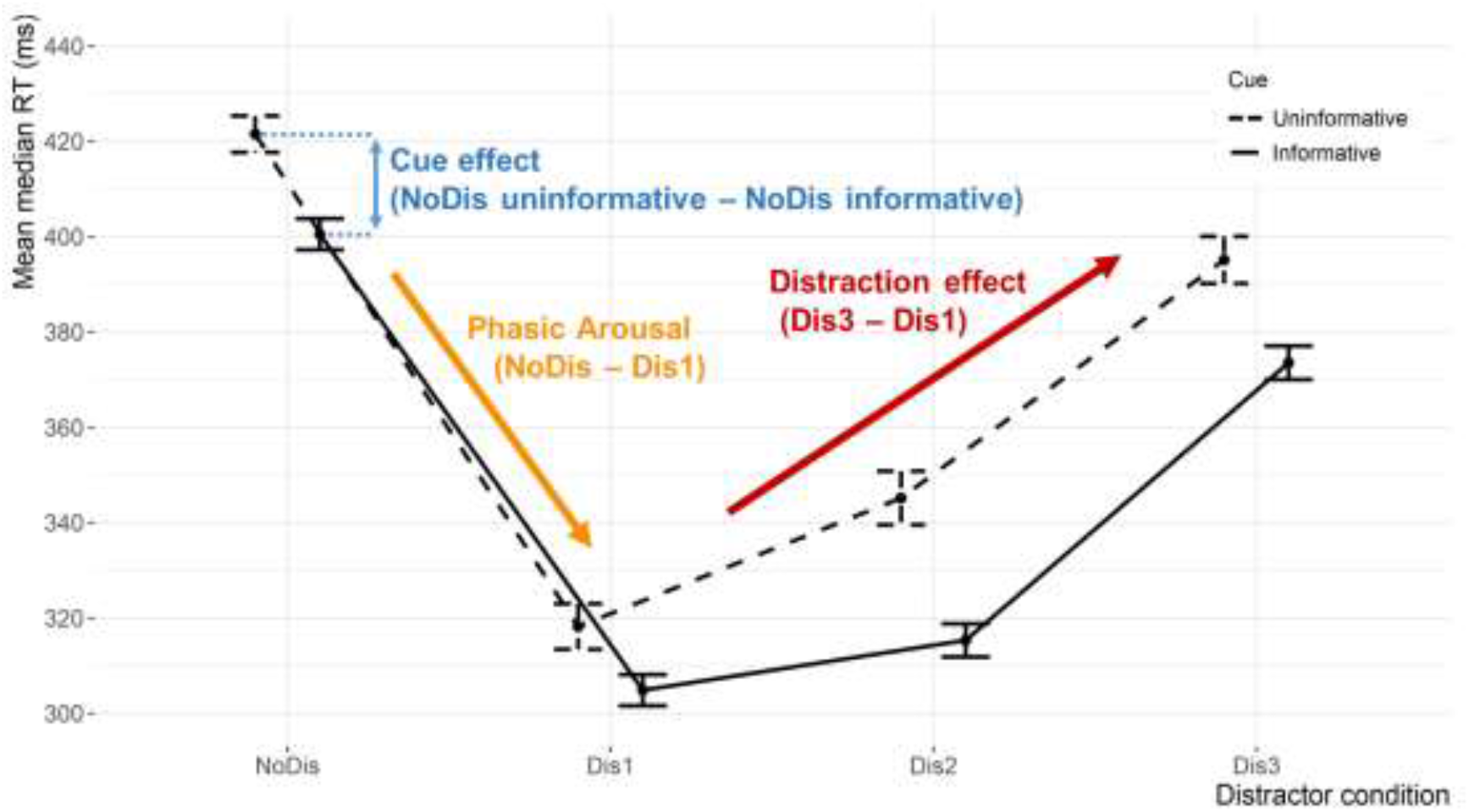
Median RT according to cue and distractor conditions. *Note*. Mean of median reaction time as a function of the cue category (informative or uninformative) and of the distractor condition (NoDis, Dis1, Dis2, Dis3). Significant differences and related p values can be found in the Median RT part in the Results section. (Error bars represent 1 SEM).

A DISTRACTOR by AGE interaction was significant (χ^2^ (39) = 81.8; p < .001). Planned post-hoc analyses were performed on the RT+ arousal facilitation and distraction effects. First, a Shapiro-Wilk test indicated that the arousal and distraction effects (see Fig. 2) were not normally distributed (W = 0.94; p < .001 and W = 0.98; p < .001, respectively). Post-hoc Kruskal-Wallis tests revealed that the AGE (χ^2^ (13) = 91.0; p < .001; Fig. 3c) had a significant effect on the arousal facilitation effect: it was larger in the 6, 7 and 8-year-olds compared to the 13 to 25-year-olds (p < .05). Other significant effects are shown in Fig 3b. This result was confirmed by decisive evidence for a negative correlation of the arousal effect normalized measure with age (Kendall’s Tau = −0.141, BF_10_ = 132.7). AGE (χ^2^ (13) = 47.4; p < .001; Fig. 3d) also had a significant effect on the distraction effect: children of 6 years of age showed higher scores than the 12 to 25-year-olds (p < .05). This was not confirmed by Bayesian statistics: a positive evidence against a correlation of the Distraction effect normalized measure with age (Kendall’s Tau = −0.044, BF_10_ = 0.1) was found.

**Figure 3.**
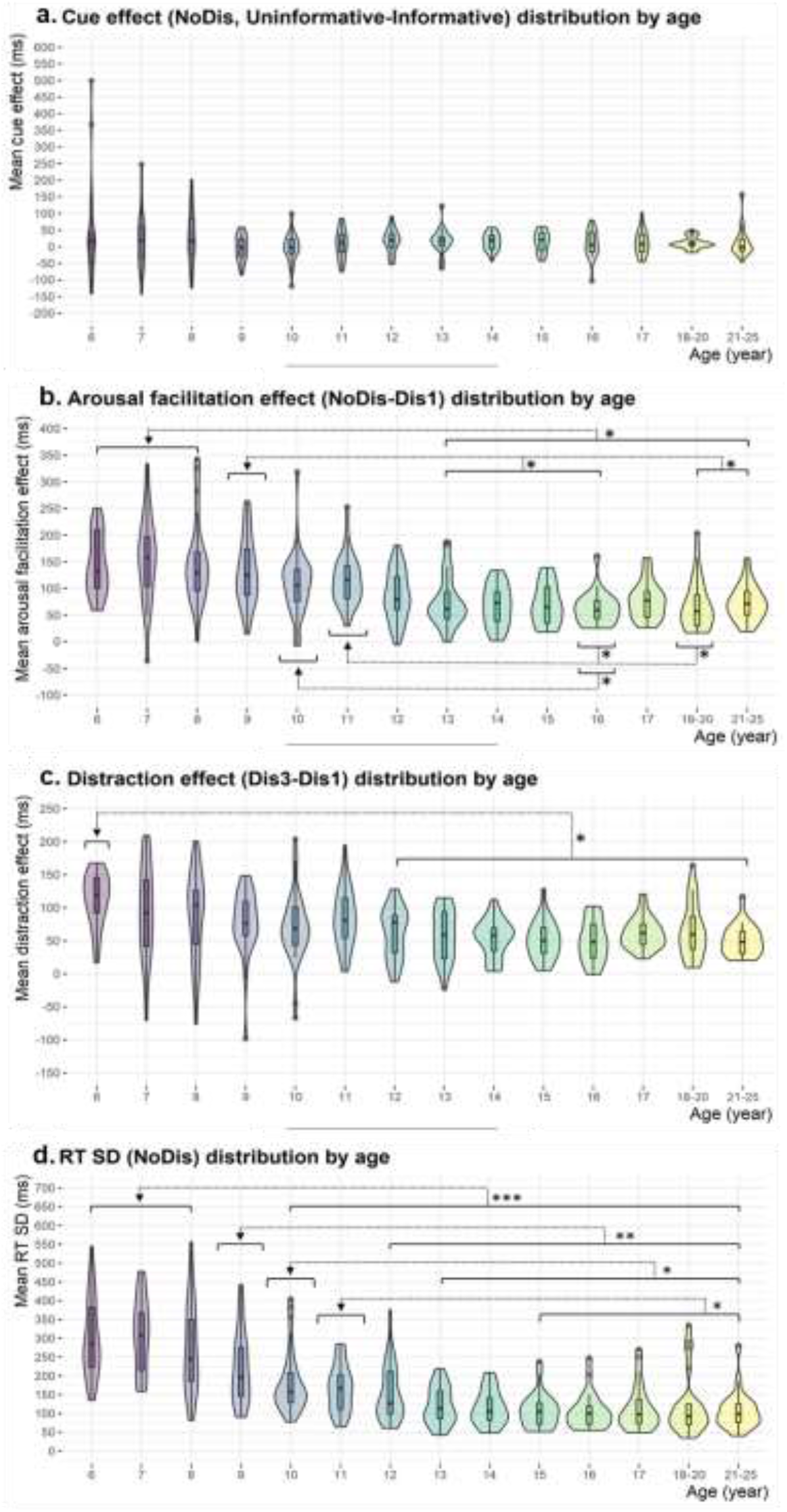
RT_+_ effects according to age. *Note*. a) Reaction time differences between NoDis uninformative and informative (cue effect) as a function of the age range. b) Reaction time differences between NoDis and Dis1 (arousal effect) as a function of the age range. c) Reaction time differences between Dis3 and Dis1 (distraction effect) as a function of the age range. d) Reaction time variability (standard deviation across blocks) as a function of age range. (p < .05 *, p < .01 **, p < .001 ***). Within each boxplot (Tukey method), the horizontal line represents the median, the box delineates the area between the first and third quartiles (interquartile range); juxtapose to each boxplot, the violin plot adds rotated kernel density plot on left and right side.

### RT SD

RT SD was modulated by AGE (χ^2^ (13) = 287.1; p < .001; Fig. 3d). HSD post-Hoc comparisons revealed that RT SD was larger in the 6 and 8-year-olds compared to the 10 to 21-25-year-olds (p < .001). Significant higher RT SD was also found in the 9-year-olds compared to the 12 to 21-25-year-olds (p < .01), in the 10-year-olds compared to the 13 to 21-25-year-olds (p < .05), and in the 11-year-olds compared to the 15 to 21-25-year-olds (p < .05). The RT SD decreases between 8 and 12-years-old.

### Response types

The distribution of the different types of responses according to age is depicted in Fig. 4. The average correct response rate was 76.0 ± 0.3%. No main effect of AGE, nor interaction with AGE, was found for CueRep (total average: 0.7 ± 0.1%; Fig. S6a) and LateRep (total average: 11.0 ± 0.2%; Fig. S6b). Significant effects of age on the other response types are detailed in the following.

**Figure 4.**
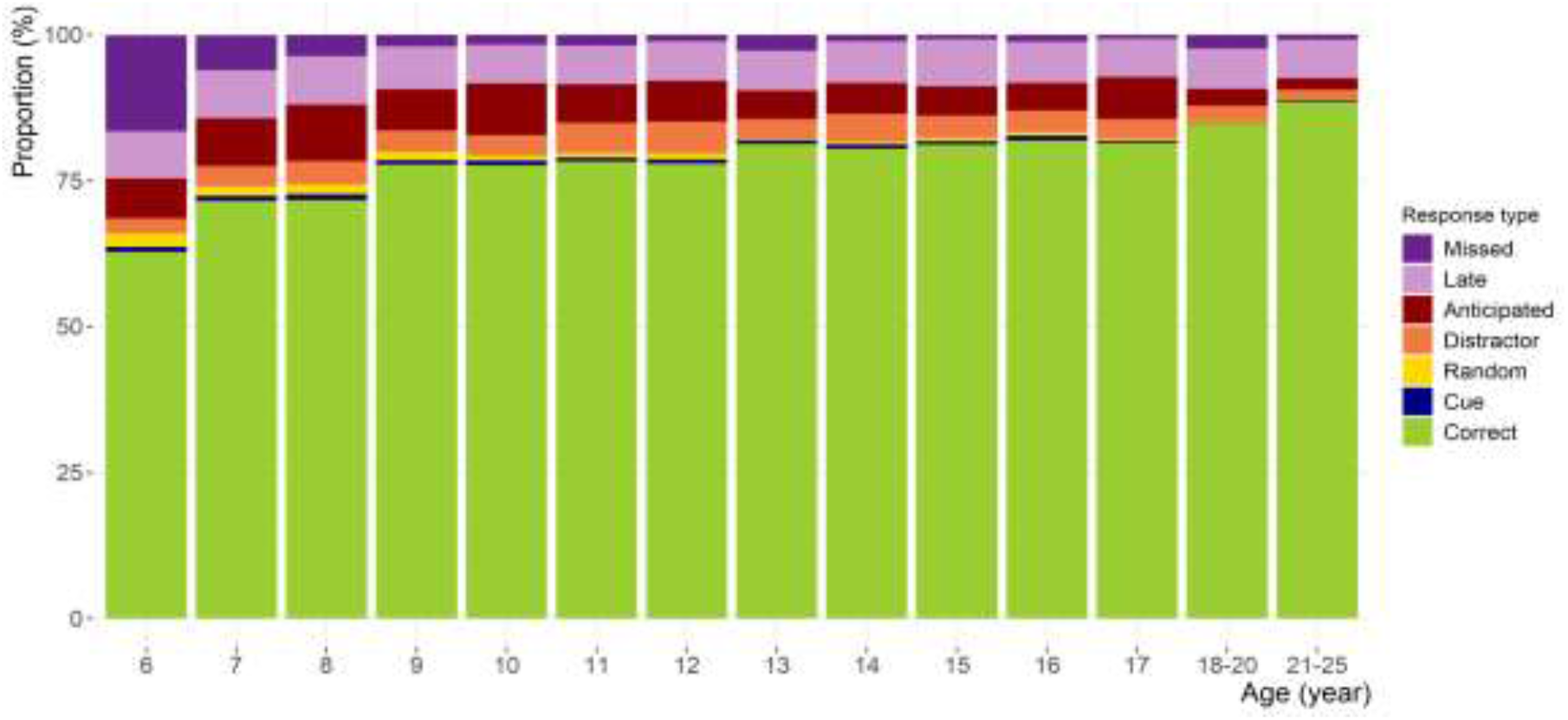
Response type proportions according to age.

### Missed responses

The rate of missed responses (3.5 ± 0,1%) was modulated by AGE (χ^2^ (13) = 96.0; p < .001) and DISTRACTOR (χ^2^ (3) = 133.8; p < .001). A DISTRACTOR by AGE interaction was also significant (χ^2^ (39) = 343.9; p < .001, Fig. 5). HSD post-hoc tests indicated significant larger MissRep rate in the Dis conditions compared to the NoDis condition in the 6 and 7-year-olds, only (p <.001). In the NoDis condition, HSD post-hoc comparisons indicated no significant difference in the MissRep rate with age (p = .124). In the distractor conditions, a higher percentage of MissRep was found in the 6 to 7-year-old children (p < .05). More precisely, the 6-year-olds had a higher MissRep rate than the 8 to 21-25-year-olds in all the distractor conditions, while the 7-year-olds presented more MissRep than (i) the 10, 12, 15, 17 and 21-25-year-olds in the Dis1 condition, (ii) the 10 and 17 to 21-25-year-olds in the Dis2 condition, and finally (iii) the 10 and 15 to 25-year-olds in the Dis3 condition. In summary, only the 6 and 7-year-olds missed target sounds preceded by a distracting sound.

**Figure 5.**
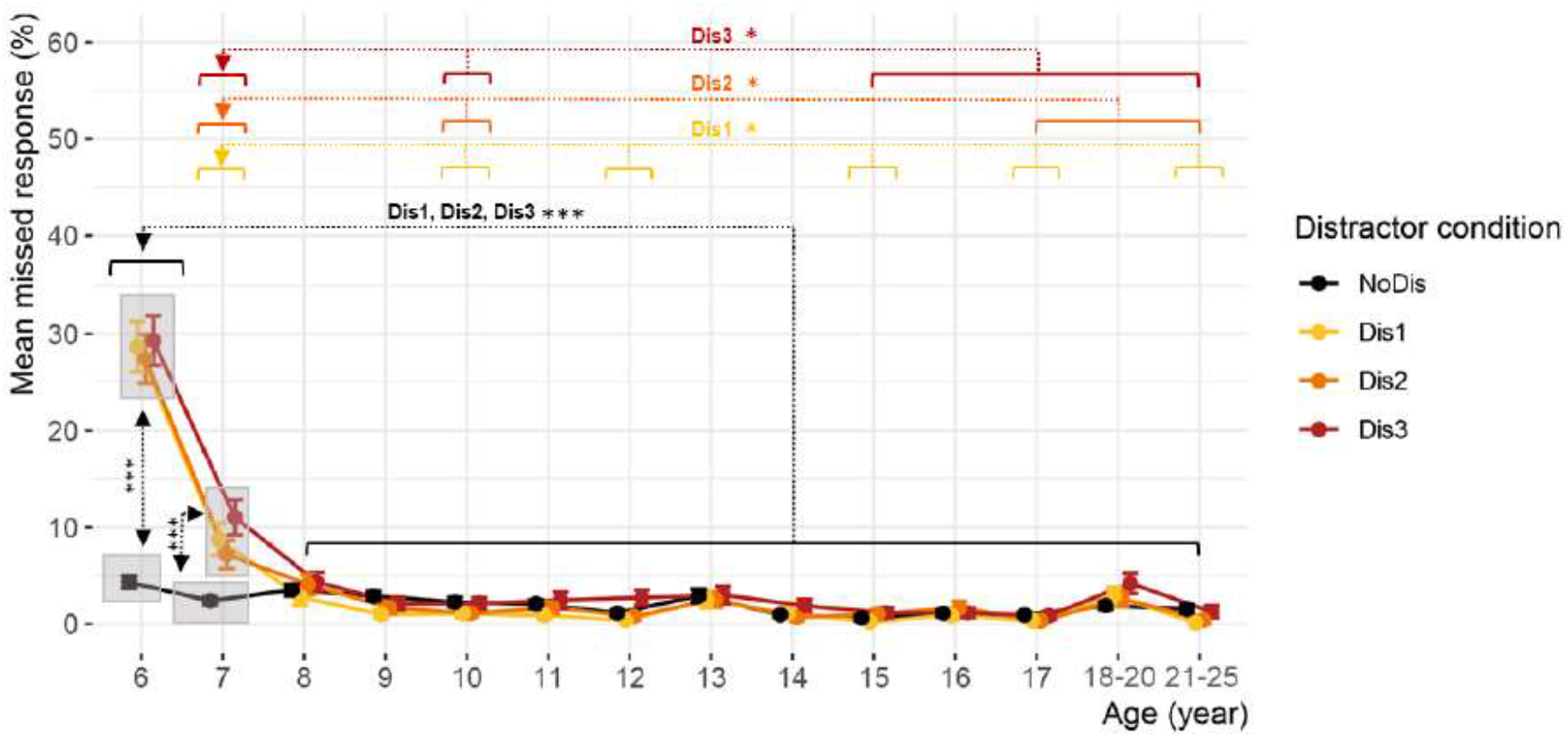
Missed responses according to age. *Note*. Mean missed responses percentage as a function of the distractor condition and the age (error bars represent 1 SEM).

### Anticipated responses

The rate of anticipated responses (10.3 ± 0.3% on average) was modulated by AGE (χ^2^ (13) = 52.9; p < .001; Fig. 6a). Post-hoc HSD analysis showed that the 7 to 12 and the 17-year-olds had more AntRep than the 21-25-year-olds (p < .05). Children from 7, 8 and 10 years of age showed an increased AntRep rate compared to the 18-20-year-olds (p < .05). Finally, the 10-year-olds showed a higher AntRep rate than the 13-year-old children (p < .05).

**Figure 6.**
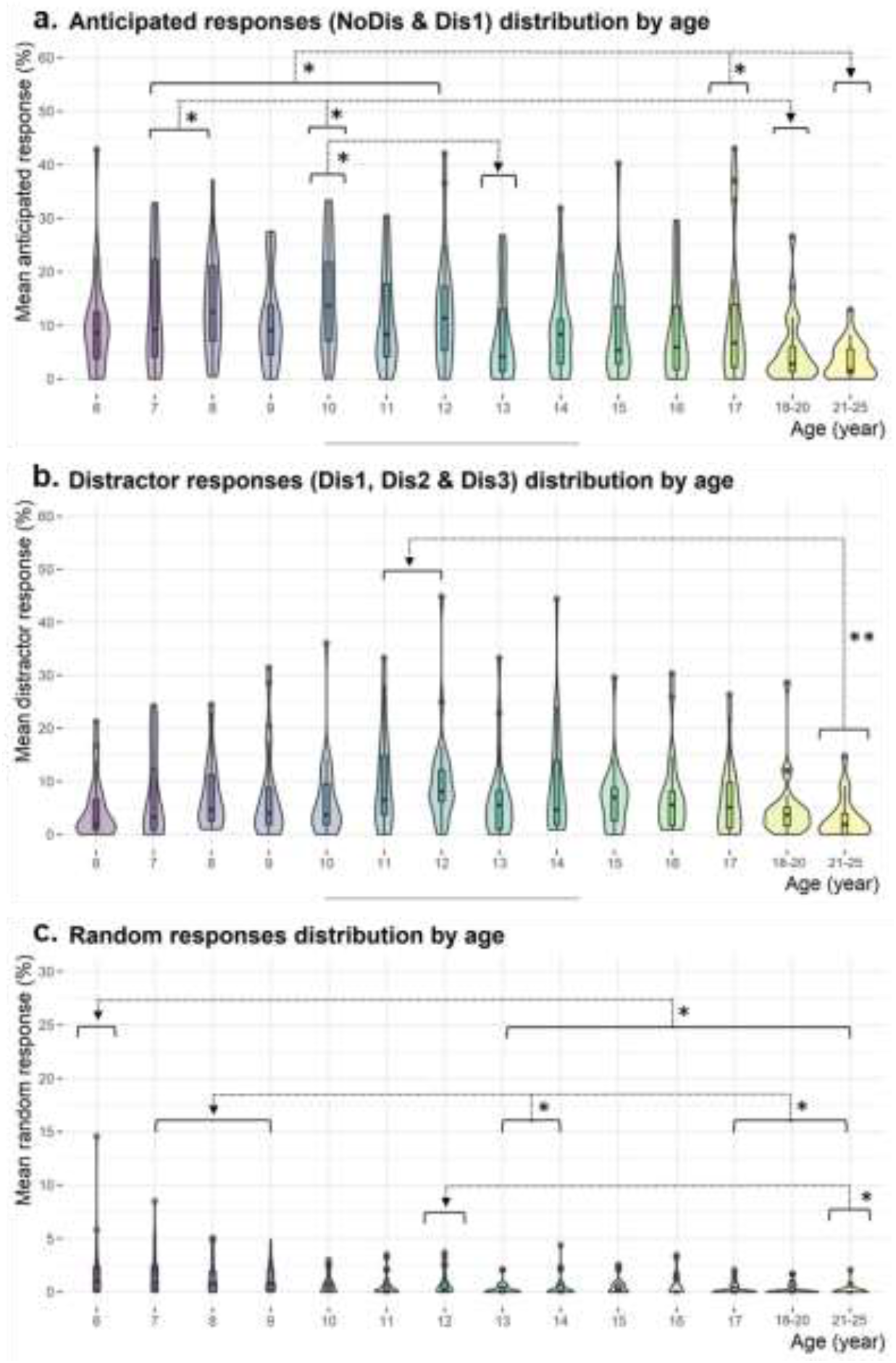
Response type proportions according to age. *Note*. a) Anticipated responses percentage (NoDis and Dis1) as function of the age. b) Distractor responses percentage as a function of the age range. c) Random responses percentage as a function of the age range. (p < .05 *, p < .01 **, p < .001 ***). Within each boxplot (Tukey method), the horizontal line represents the median, the box delineates the area between the first and third quartiles (interquartile range); juxtapose to each boxplot, the violin plot adds rotated kernel density plot on each side.

We also observed a significant effect of GENDER (χ^2^ (1) = 10.3; p = .001) indicating larger AntRep rate in male (11.7 ± 0.4%) compared to female (8.9 ± 0.4%) participants.

We observed significant main effects of CUE (χ^2^ (1) = 18.7; p < .001) and DISTRACTOR (χ^2^ (1) = 702.6; p < .001), as well as a significant DISTRACTOR by CUE interaction (χ^2^ (1) = 15.3; p < .001). According to HSD post-hoc comparisons, participants made more AntRep in the Dis1 (left: 21.2 ± 0.9% / right: 17.2 ± 0.8% / neutral: 18.3 ± 0.8%) than in the NoDis (left: 2.2 ± 0.2% / right: 2.2 ± 0.2% / neutral: 1.4 ± 0.2%) condition (p < .001), independently of the cue nature. The AntRep rate was found larger with informative cues rather than with uninformative ones in the NoDis condition (both left and right informative cues: p < .001); while it was greater with left cues compared to right and neutral cues in the Dis1 condition (both: p < .001).

### Distractor responses

The rate of distractor responses (7.0 ± 0.2% on average) was found modulated by AGE (χ^2^ (13) = 30.8; p = .004; Fig. 6b): the 11 (9.7 ± 0.8%) and 12 (10.0 ± 0.8 %) year-old children made more DisRep than the 21-25-year-olds (3.2 ± 0.5%; p < .01).

We also observed a significant main CUELRN (χ^2^ (13) = 48.5; p < .001) effect: all participants made more Disrep in the left cue condition (8.8 ± 0.3%) than in the right (7.0 ± 0.3%; p < .001) and the neutral ones (6.1 ± 0.3%; p < .001).

### Random responses

The rate of random responses (0.8 ± 0.1% on average) was modulated by AGE (χ^2^ (13) = (77.2); p < .001; Fig. 6c). According to HSD post-hoc comparisons, the 6-year-olds (2.0 ± 0.3%) made more RandRep than the 13 (0.3 ± 0.1%), 14 (0.5 ± 0.2%), 15 (0.6 ± 0.2%), 16 (0.5 ± 0.2%), 17 (0.3 ± 0.1%), 18–19 (0.3 ± 0.1%) and 21-25 (0.1 ± 0.1%) year-olds (p < .05). The 7 (1.8 ± 0.3%), 8 (1.4 ± 0.2%) and 9 (1.2 ± 0.2%) year-olds made more RandRep than both the 13 and 14-year-olds, and the 17 to 25-year-olds (p < .05). Additionally, the 12-year-olds (0.8 ± 0.1%) made more RandRep than the 21-25-year-olds (p < .05).

The percentage of incorrect responses decreases with age. Incorrect responses are due to distracting sounds inducing a large number of missed responses in the youngest ones (age 6 and 7) and a great amount of responses to distractors in the 11 and 12-year-olds. Moreover, the 6-9-year-olds present a higher rate of random responses and the 7-12-year-olds a greater rate of anticipated responses (see Fig. 6 for a graphical representation of the main results according to age).

## Discussion

We aimed to characterize the developmental trajectories of attentional and motor processes related to distractibility using several behavioral measures (see a graphical summary in Fig. 7). Our findings suggest that voluntary orienting of attention is mature at 6 years of age, while voluntary sustained attention slowly develops until the age of 12. Distraction is greater before the age of 8, compared to older age groups. Later in childhood and adolescence, there is increased impulsivity, which fades in adulthood.

**Figure 7.**
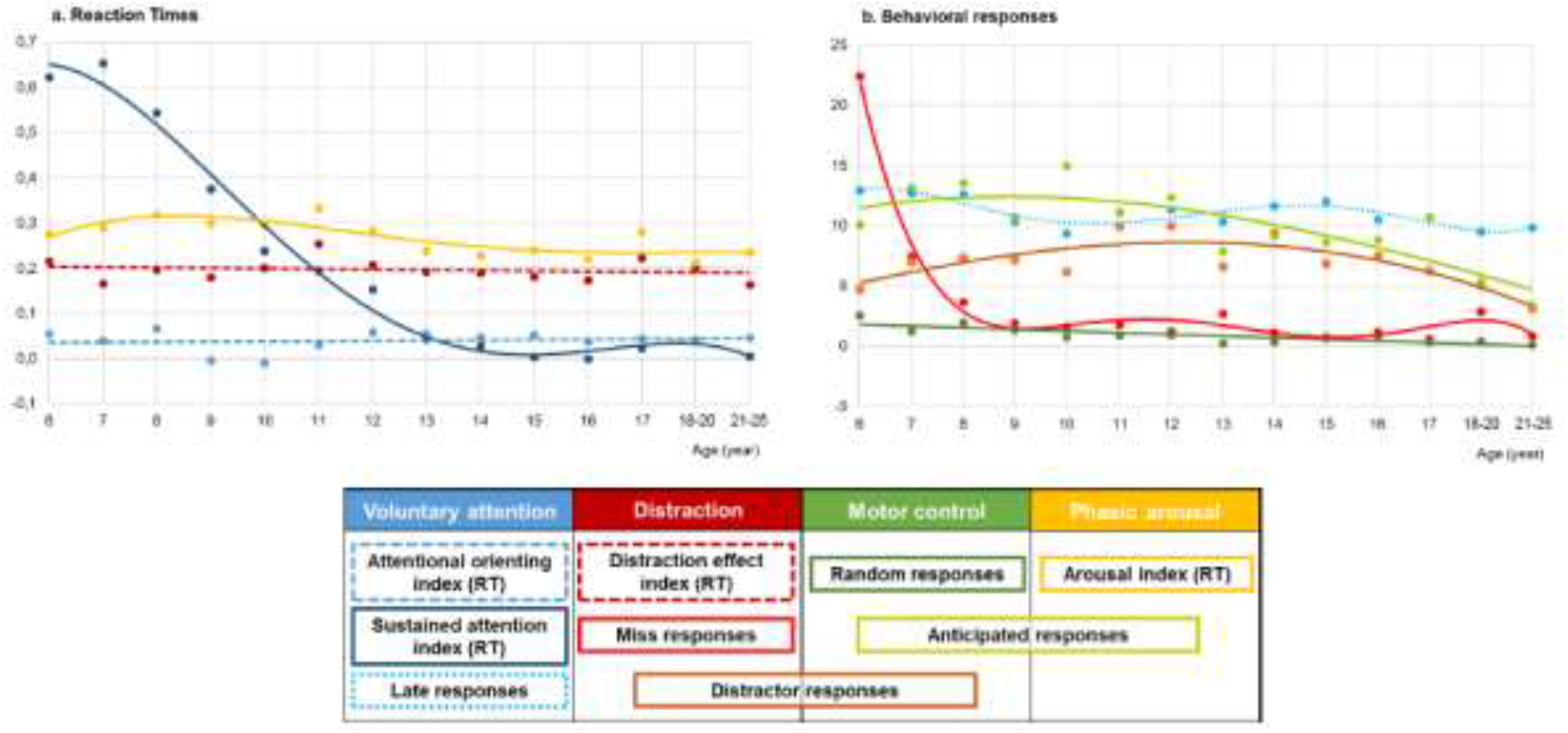
Graphical representation of the main results. *Note*. a) Reaction times normalized measures according to age. Curves correspond to polynomial fitting curves for the Sustained Attention (order 4) and Arousal (order 4) measures, and to fitting lines for the Distraction and Attention Orienting measures. Sustained attention measure = mean RT SD for each age range normalized across age ranges; Attention orienting measure = (medianRT_+NoDisUninf_ – medianRT_+NoDisInf_) / medianRT_+AllConditions;_ Arousal measure = (medianRT_+NoDis_ – medianRT_+Dis1_) / medianRT_+AllConditions;_ Distraction effect measure = (medianRT_+Dis3_ – medianRT_+Dis1_) / medianRT_+AllConditions_. b) Percentage of late responses (LateRep), miss responses (MissRep), responses to distractors (DisRep), anticipated responses (AntRep) and random responses (RandRep) according to age. Dots represent the mean percentage. Curves correspond to polynomial fitting curves for LateRep (order 5), MissRep (order 5) DisRep (order 3) and AntRep (order 3), and to a fitting line for RanRep. Measures reflecting (1) voluntary attention are in blue colors, (2) distraction are in red colors; (3) motor control are in green colors, and (4) arousal are in yellow color. Brown and light green colors represent processes overlaps. Dotted lines represent measures which have not been found modulated by age.

Global scores from the ADHD-RS clearly indicate that parents report better attention capacities in older (12 to 17 years old) than in younger (6 to 10 years old) children. This result, consistent with previous findings (Petton et al., 2019), confirms an ongoing maturation of attentional processes and motor control during childhood and adolescence.

### Voluntary attention capacities in children

Shorter reaction times to targets preceded by informative rather than uninformative cues is typically used as a measure of voluntary attention orientation towards the cued side in the informative condition (Posner, 1980, 2012). According to Bayesian correlation analysis performed in the current study, there is no evidence for an effect of age on this cue effect in the absence of distracting sounds, corroborating previous studies showing mature voluntary attention orienting at 6 years of age, or even during the first year of life (Ross-Sheehy et al., 2015; Rueda et al., 2004).

In the absence of distracting sounds, difficulties in sustained attention can result in increased RT variability and late response rate (index of attentional lapses). Increased intra-subject variability of RT reflects voluntary attentional efficiency (Antonini et al., 2013). We found that RT variability between trials with no distracting sounds during the whole experiment (around 15 min) slowly decreases between the ages of 8 and 12 corroborating an improvement in sustained attention during this period (Lin et al., 1999; Petton et al., 2019; Thillay et al., 2015). This progressive maturation of sustained attention between the ages of 8 and 12 may be related to the structural and functional maturation of the frontal lobes (Blakemore & Choudhury, 2006; Gogtay et al., 2004), allowing a more efficient voluntary attentional control (Blakemore & Choudhury, 2006; Thillay et al., 2015). Interestingly, we did not find an increase in RT variability through the task as previously observed in adults (Flehmig et al., 2007). This absence of modulation over time could be explained by the presence of distracting sounds in 50% of the trials in the CAT, compared to typical paradigms used to measure sustained attention such as the Continuous Performance Test (Kanaka et al., 2008; Lin et al., 1999). Distractors trigger a phasic arousal burst (Bidet-Caulet et al., 2015), which increases alertness for a few seconds (Aston-Jones & Cohen, 2005). This could help in maintaining an appropriate general arousal level compensating the vigilance decline across the blocks. Even when selecting the trials without distractors to analyze sustained attention, phasic arousal could still have an effect, especially when the trials without distractors were preceded by a late occurrence distractor trial (Masson & Bidet-Caulet, 2019). We also found no evidence for an effect of age on the rate of late responses reflecting attentional lapses, in contrast with previous studies highlighting a global decrease in spontaneous fluctuation of attention between 8 and 12 year of age (Petton et al., 2019). Phasic arousal could also partially compensate for decreased sustained attention capacities by reducing RT variability enough to avoid attentional lapses. Therefore, the CAT seems to be valuable to measure the efficacy of sustained attention in a context with distractors.

These findings further suggest that some proactive form of attentional control is already present at the age of 6 to improve anticipation and detection of a target in accordance with previous information, but that it still needs to mature to achieve stabilized performances over time.

### Impact of distracting information on children behavior

Several measures of the CAT reflect distraction (i.e. a behavioral cost). Distracting sounds can result in longer reaction times (Bidet-Caulet et al., 2015; ElShafei, Fornoni, et al., 2018; Masson & Bidet-Caulet, 2019), or worse, in missed responses to the following target (Parmentier, 2008; Parmentier & Andrés, 2010) and sometimes to responses to the distractor (van den Wildenberg & Crone, 2005). The strength of the CAT lies in the different timings of the distractor sounds before the target, enabling the dissociation of the behavioral cost and benefit they induce. In line with previous studies using the CAT in adults (Bidet-Caulet et al., 2015; ElShafei, Bouet, et al., 2018; ElShafei et al., 2019; Masson & Bidet-Caulet, 2019), we observed two distinct effects on RTs triggered by the distracting sounds. First, distracting sounds played long before the target (Dis1 and Dis2) induced a reduction in reaction times compared to a condition without distractor (NoDis). This benefit in RT has been attributed to an increase in phasic arousal (Masson & Bidet-Caulet, 2019), but the contribution of a warning cue effect of the distracting sound cannot be ruled out. Second, distracting sounds played just before the target (Dis3) resulted in an increase in reaction times compared to conditions with a distractor played earlier (Dis1 and Dis2). This cost in RT is considered as a good behavioral approximation of distraction (Bidet-Caulet et al., 2015; ElShafei et al., 2019; Masson & Bidet-Caulet, 2019), though a longer foreperiod before targets preceded by early rather than late distractors can also contribute to this effect (e.g. Drazin, 1961; Niemi & Näätänen, 1981). Importantly, this measure of distraction using the CAT (difference ifn reaction time to targets preceded by the same distracting sound at different timings) is slightly different from the deviance distraction effect (difference in reaction time to targets preceded by or including a standard vs a deviant distracting event) obtained with oddball paradigms. Indeed, the CAT rather provides a measure of distraction by an unexpected isolated salient event; while oddball paradigms provide a measure of distraction triggered by deviants embedded in a regular sound sequence. In audiovisual oddball paradigms, the sound sequence (including the deviants) could be inhibited by proactive selective attention mechanisms, similar to the unattended stream in a classic dichotic paradigm (Bidet-Caulet et al., 2010; Hillyard et al., 1973); while in auditory-auditory oddball paradigms, the processing of distracting and relevant information, both embedded in the same stream, can interfere. This is less likely to happen with the CAT since distractors are clearly dissociated from the relevant information and not embedded in a regular stream of sounds.

Both phasic arousal and distraction effects were observed between the ages of 6 and 25. The CAT thus enables the dissociation of the effects of arousal (RT NoDis – Dis1) and distraction (RT Dis3 – Dis1) on the RT in both adults and children. Importantly, the developmental trajectories of these two measures were found to be different. Distraction is increased in 6-years-old only, and progressively decreases from the age of 7 to 12, although this effect is not seen when normalizing using the median RT across all trials. Phasic arousal is stable between the ages of 6 and 9, decreases between the ages of 9 and 13 and reaches the adult developmental level at the age of 13. To our knowledge, there is no literature on the developmental trajectory of phasic arousal, but a study showing increased pupil dilation to rare complex sounds in infants compared to adults (Wetzel, Buttelmann, et al., 2016). Concerning distraction, previous developmental studies using oddball paradigms provided contradictory results regarding changes in the deviance distraction effect between children and adults. Some studies found that the deviance distraction effect decreases with age, with mature performance around the age of 10 (Wetzel et al., 2006, 2018; Wetzel, Schröger, et al., 2016; Wetzel & Schröger, 2007), while other works found no effect of age (Horváth et al., 2009; Leiva et al., 2016; Ruhnau et al., 2013; Wetzel et al., 2009). This discrepancy can result from differences in the tested age-ranges and from the dissociation between the distracting event and the target. Indeed, in audiovisual oddball paradigms, distraction effect reflects the impact of a distracting sound on the response to a visual target, while in the auditory oddball paradigms, the distracting event is a feature of the target sound to discriminate. Attentional mechanisms might be differently activated in these two kinds of paradigm, in particular inhibitory mechanisms suppressing distractor processing might be less recruited in the auditory oddball paradigm. The final impact of distracting events on the performance of an unrelated task is contingent to the strength of the distraction effect and on the level of phasic arousal. Phasic arousal depends on the sound properties (Max et al., 2015b; Parmentier & Andrés, 2010; SanMiguel et al., 2010; Wetzel et al., 2012) and the level of tonic arousal (Aston-Jones & Cohen, 2005; Howells et al., 2012) which are influenced by the task demands (Eysenck, 1982; Kahneman, 1973; Yerkes & Dodson, 1908). In future research, it seems important to measure levels of tonic and phasic arousal to understand the developmental trajectory of distraction. The present results suggest that phasic arousal and distraction would follow distinct developmental trajectories. However, the precise relation between the arousal and distraction effects still needs to be characterized and the CAT measures should not be conceived as makers of independent cognitive functions, but as complementary measures of the impact of distractors on performance.

The number of missed and incorrect responses in attentional tasks is a sensitive measure of distraction since it was found to negatively correlate with school performance (Zimmermann & Fimm, 2002). In the presence of distracting sounds (irrespective of their timing), we observed a large increase of missed responses only in the 6 and 7-year-old children. This detrimental distraction effect strongly decreases from the age of 6 to 7, and moderately from the age of 7 to 8. At 8-years-old, the missed response rate reaches the adult level (with or without distracting sounds). An increase in missed response rate was previously seen in children of 5 and 6 years of age in a no-distractor context (Kanaka et al., 2008). In contrast, the missed response rate was not modulated by age in the CAT no-distractor trials, suggesting that the missed responses in the 6 and 7-year-olds is caused by the deleterious effect of the distracting sound. Additionally, the 11 and 12-year-old children responded more to distractors than the 20 to 25-year-olds. This increase in responses to the distractor suggests a higher impulsivity at this age, which progressively decreases from 13 to 19 years of age. A decrease in responses to irrelevant stimuli from the ages of 3 to 16 was previously observed (Booth et al., 2003; Ridderinkhof et al., 1999; van den Wildenberg & Crone, 2005; Wright et al., 2003). Taken together, these results suggest that resistance to interference improves during childhood until late adolescence.

In the CAT, longer reaction times, target omission and responses to a distractor all indicate the predominance of the reactive form of attentional control in young children. They could result from either (i) an increased involuntary attentional capture, (ii) a reduced voluntary attentional inhibition of distracting sounds, or finally, (iii) an impossibility/difficulty to reorient the attentional focus back to the task at hand. Until now, few studies have investigated these hypotheses. Some electroencephalographic and behavioral works suggest that the increased behavioral distraction in children results from a delayed reorientation of attention to the task at hand (Ruhnau et al., 2010; Wetzel et al., 2018). However, inconsistent results have been reported (Wetzel & Schröger, 2014) and further electro- or magnetoencephalographic studies during development will help our understanding of the brain mechanisms underlying increased distractibility.

In summary, distraction is increased in the youngest children (ages 6 and 7) reflected by a large increase of missed responses. In the 11 and 12-year-olds, enhanced distraction manifested as increased impulsivity, reflected in an increase in responses to distracting sounds. The combined use of reaction times, as well as missed and distractor response rates is necessary to assess the developmental trajectory of distraction and phasic arousal triggered by distracting sounds. This will help to fully understand the impact of distractors on behavior. Distraction is multifaceted and results in both attention and motor manifestations.

### Beyond attention: Impulsivity and motor control in children

In broader models of behavioral control, the executive system coordinates the interaction of memory, attention and motor processes (Diamond, 2013; Kahneman, 1973). Motor control and attention are tightly linked: motor inhibition is driven by attentional selection, which is conditioned by past actions and their related memory traces. Difficulties in motor inhibition can result in responses to task-irrelevant events such as the distracting sounds or responses in anticipation of the targets, which can be considered as the behavioral expression of impulsivity. Many models have suggested a relation between enhanced arousal level, impulsivity and motor control (e.g., Barratt & Patton, 1983; Eysenck & Eysenck, 1985). While the development of phasic arousal is poorly documented, impulsivity and motor control were found to be enhanced and reduced, respectively, in children (Booth et al., 2003; Kanaka et al., 2008; Ridderinkhof et al., 1999; van den Wildenberg & Crone, 2005; Wright et al., 2003).

While quite variable with age, the rate of anticipated responses is relatively stable between the ages of 7 and 12 and between the ages of 13 and 17. It first decreases around 12-13 years of age, and then again around 17-18 years of age. Both an inaccurate prediction of the target time occurrence and difficulties in voluntarily maintaining motor control could account for anticipated responses. Higher anticipated response rates might reflect an immature proactive strategy progressively developing from childhood to adulthood. Increased impulsivity in children before the age of 12 has previously been observed (Kanaka et al., 2008; Lin et al., 1999; Thomas et al., 1981). Participants made anticipated responses to the target mostly in the presence of a distractor irrespective of age, suggesting that processes triggered by distractors influence the behavioral expression of anticipated responses. These anticipatory responses following distracting sounds could be driven by the phasic increase in arousal triggered by distractors or by reduced voluntary inhibitory motor processes.

We also noticed a progressive decrease in random response rate between the ages of 10 and 12. As random response timing corresponds to a response which is believed to be independent from a stimulus, this response would be related to motor – rather than attentional – control. Our findings suggest that motor control is mature enough at the age of 13 to reach adult performance level.

### Conclusion

Despite its intrinsic limitations (e.g. testing done on different experimental sites, stress and motivation levels of the participants not considered in the analyses, no measurement provided for tonic arousal, scarcity of some responses when crossing experimental conditions), this study provides critical and precise information about the developmental trajectories of several facets of distractibility from childhood to adulthood.

Voluntary orienting is found to be functional at 6 years of age, while sustained attention gradually develops from the ages of 8 to 12. Interestingly, distraction manifests as omissions of relevant stimuli in 6-7-year-olds, as impulsivity in 11-12-year-olds, and as delayed reaction times to relevant stimuli from 6 years of age to adulthood. These findings suggest that the attentional imbalance resulting in increased distractibility is rather related to reduced voluntary sustained attention capacities and enhanced distraction in children (6-8 years old), but to decreased motor control and increased impulsivity in teenagers (10-17 years old).

Taken together, these findings are in line with a reactive to proactive shift in attention and motor control during childhood (Blackwell & Munakata, 2014; Chevalier et al., 2014; Doebel et al., 2017; Munakata et al., 2012). Interestingly, if the reactive form is predominant during early childhood, the present results show that some proactive control is already present at 6 years of age and progressively matures until young adulthood.

In light of the present findings, psycho-education and classroom learning strategies could be improved by targeting attention processes in children and motor control capacities in young teenagers, as of now. As few standardized neuropsychological tests are currently available to assess distractibility, assessing the validity of the measures used in the present study could help to better characterize attentional deficits and improve individualized care for patients.

## Acknowledgements

We thank P.R. Bauer and R. Masson for their careful reading of this manuscript. We also thank O. Abdoun for his helpful advices concerning the statistical approach used in this study. We wish to thank all our partner in the education field, especially E. Subra, for their help in recruiting subjects. Eventually, we would like to thank all the participants and their parents for their time.

This work was performed within the framework of the LABEX CORTEX (ANR-11-LABX-0042) and the LABEX CeLyA (ANR-10-LABX-0060) of Université de Lyon, within the program “Investissements d’Avenir” (ANR-16-IDEX-0005) operated by the French ANR. The CAT is protected by a pending trademark application « CoLeT ».

## Supporting information

**Figure S1.** Repartition of socio-economical statuses across the population sample.

**Figure S2.** Lower RT limit.

**Figure S3.** Schematic view of the cost and benefit triggered by distracting sounds.

**Figure S4.** Timeline for the behavioral response categorization during the CAT trials.

**Figure S5.** Median RT according to the age.

**Figure S6**. Behavioral responses according to age.

**Table S1.** Results of Bayesian repeated-measure ANOVA on auditory thresholds.

**Appendix S1.** Instructions.

**Appendix S2.** RT lower limit calculation.

**Appendix S3.** Statistical analysis of auditory thresholds.

**Appendix S4.** Statistical analysis of attention scores.

